# Expanding the *C. elegans* toolkit with gonad explants

**DOI:** 10.64898/2026.03.26.714430

**Authors:** Réda M. Zellag, Eric Cheng, Abigail R. Gerhold, Jean-Claude Labbé

**Affiliations:** Department of Biology, McGill University, 1205 avenue Docteur Penfield, Montréal, QC, H2A 1B1, Canada; Institute for Research in Immunology and Cancer (IRIC), Université de Montréal, C.P. 6128, Succ. Centre-ville, Montréal, QC, H3C 3J7, Canada; Department of Pathology and Cell Biology, Université de Montréal, C.P. 6128, Succ. Centre-ville, Montréal, QC, H3C 3J7, Canada

**Author notes:** Corresponding authors: Abigail Gerhold and Jean-Claude Labbé.

**Keywords:** *Caenorhabditis elegans*, gonad development, mitosis and meiosis, tissue explant, *ex vivo* culture, drug treatment

## Abstract

Animal development is a complex process that requires the coordination of a plethora of pathways in space and time. In several species, the availability of tissue explants has provided a simplified context that facilitates mechanistic investigations, particularly into dynamic events. Here, we demonstrate that extruded *C. elegans* gonads are a viable tissue explant system for this model organism. Using live-cell imaging, we show that *C. elegans* gonad explants retain many tissue properties that have been documented *in vivo*, including mitosis, meiosis, apoptosis and gametogenesis. We further show that *C. elegans* explants are acutely responsive to treatment by the microtubule depolymerizing drug nocodazole. Our work thus reveals *C. elegans* gonad explants as a new system in which live-cell imaging and acute drug treatment can be combined to decipher the mechanisms governing germline development.

## Introduction

The coordination of cellular behaviours enables proper cell fate specification and tissue morphogenesis during development. While *in vivo* genetic manipulations have provided critical insight into the molecular mechanisms that govern this coordination, they often lack the temporal resolution needed to dissect dynamic interactions and can be complicated by indirect and compounding effects. Tissue explants, and more recently organoids, offer a complementary *ex vivo* approach that allows dynamic cellular events to be examined in complex, intact tissues (Liberali and Schier, 2024; Mogollón and Ahtiainen, 2020).

The *C. elegans* gonad is a widely used model organ that has provided insight into a variety of developmental processes, including niche-stem cell interactions, stem cell biology, meiosis, gametogenesis, sex determination, apoptosis, germline-soma interactions and tissue morphogenesis (Gartner et al., 2008; Hillers et al., 2017; Hubbard and Schedl, 2019; Huelgas-Morales and Greenstein, 2018; Joshi et al., 2010; Korta and Hubbard, 2010; Meyer, 2022; Tolkin and Hubbard, 2021; Wang et al., 2022; Wong and Schwarzbauer, 2012; Zellag et al., 2025b). Although both classical and sophisticated genetic approaches have been used to great effect in this system, a robust explant-based method for studying gonad development is currently lacking.

In *C. elegans* adult hermaphrodites, the gonad consists of two U-shaped arms that connect to the ventral uterus at their proximal end (reviewed in (Hubbard and Schedl, 2019)). The distal end of each arm is capped by a somatic distal tip cell (DTC) that acts as niche for the underlying germ cells, maintaining their mitotic competency. Germ cells that exit the niche progress through meiosis and differentiate into mature gametes at the proximal end. In late larvae, the most proximal germ cells differentiate to form a fixed number of sperm, after which germ cells produce only oocytes for the remainder of the animal’s life. Germ cells are also influenced by other somatic cells, including the sheath cells that enwrap large regions of each gonad arm, and the intestine that is located adjacent to the gonad (Hall et al., 1999; Killian and Hubbard, 2005; McCarter et al., 1997).

We recently developed a method to cultivate extruded *C. elegans* gonads *ex vivo* as explants and used it to show that germ cell spindle orientation is regulated gonad autonomously (Zellag et al., 2025a). Here, we expand this method to demonstrate that many aspects of gonad development are preserved in gonad explants. We further show that gonad explants are amenable to acute perturbation by the microtubule depolymerizing drug nocodazole. Gonad explants thus expand the toolkit for deciphering mechanisms of *C. elegans* germline development and pave the way for small molecule inhibitor-based studies to acutely perturb gene activity in this tissue.

## Results and Discussion

### Extruded *C. elegans* gonads maintain key developmental processes

To obtain viable *C. elegans* gonad explants, we took advantage of published methods for dissecting gonads for immunofluorescence to extrude intact gonad arms containing germ cells, the DTC, and sheath cells, enclosed within the gonad basement membrane (Ananthaswamy et al., 2022), into media that was developed for *ex vivo* culture of *C. elegans* embryonic blastomeres (so-called “meiosis media” (Edgar and Goldstein, 2012); Figure 1A). We found that most extruded gonads maintained their normal morphology and appeared viable for at least two hours of culturing. This was consistent for gonads from developing (L3 and L4 larval) and mature (1-day old adult) hermaphrodites, as well as from males at the late larval stage (Figure 1B-E; Movie 1-3).

**Figure 1.**
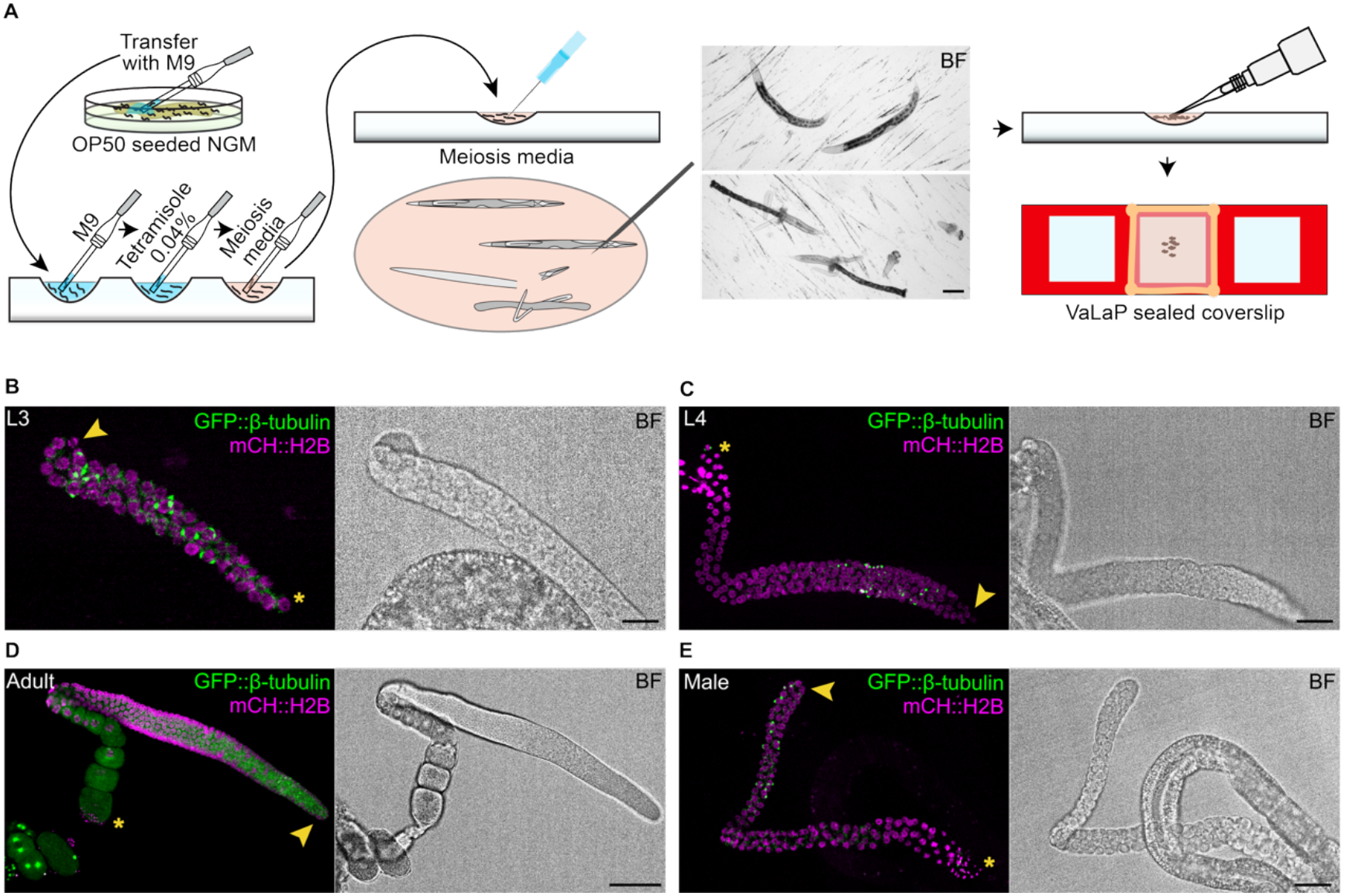
Generation of *C. elegans* gonad explants. (A) Schematic illustrating the method for obtaining gonad explants, from worm collection (left), dissection (middle) and mounting (right). The bright field (BF) images show animals before (top) and after (bottom) dissection. Scale bar = 100 µm. (B-E) Maximum intensity projections of GFP::β-tubulin (green) and mCH::H2B (magenta) (left) and bright field images (right) from gonad arms extruded from hermaphrodites at the L3 (B), L4 (C) and adult (D) stages, and a male at the L4 stage (E). Asterisks and arrowheads indicate the proximal and distal ends of the gonad, respectively. Scale bars = 10 µm (B), 20 µm (C, E), 50 µm (D).

We used live-cell imaging of GFP-tagged TBB-2 (hereafter GFP::β-tubulin) to measure the duration of germ cell mitosis (nuclear envelope breakdown to anaphase onset) and found no difference between germ cells in explants compared to those in intact animals (Figure 2A-B; Movie 4). From this we infer that germ cell physiology is not overtly perturbed in explants, as defects in germ cell spindle assembly activate the spindle assembly checkpoint (SAC) and delay mitotic exit (Gerhold et al., 2015; Zellag et al., 2021).

**Figure 2.**
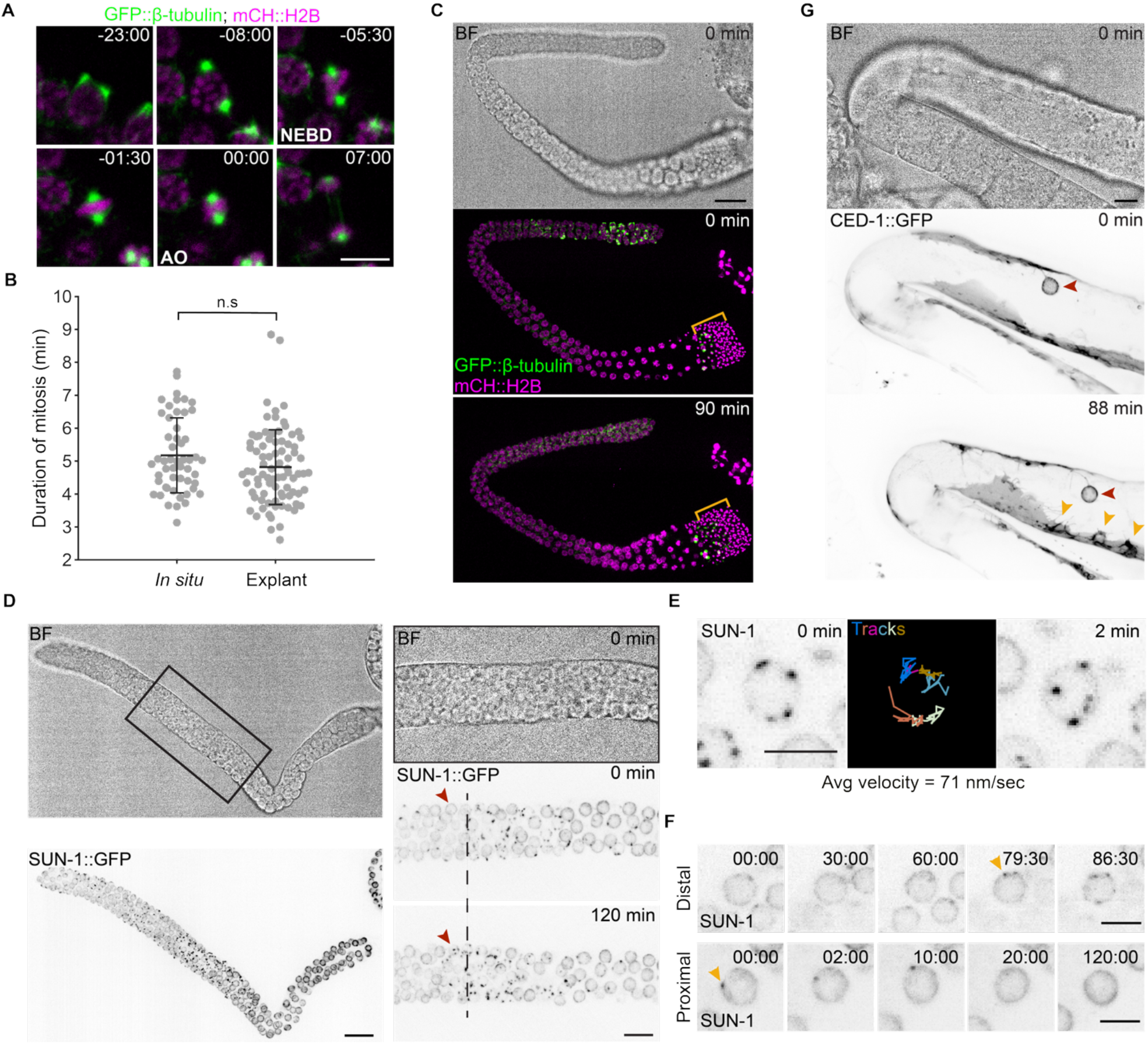
*C. elegans* gonad explants can be used to visualize mitotic, meiotic and apoptotic dynamics. (A) Cropped timelapse images of an explant germ cell undergoing mitosis. The spindle is marked by GFP::β-tubulin (green) and chromosomes by mCH::H2B (magenta). Numbers indicate time in minutes:seconds relative to anaphase onset (AO). Scale bar = 5 µm. (B) Duration of mitosis (NEBD to AO) in germ cells *in situ* and in gonad explants. Bars indicate average ± standard deviation. Each dot represents one germ cell. n.s. = not significant (p=0.0789) by a two-sample t-test. (C) Bright field image (top) and maximum intensity projections of GFP::β-tubulin (green) and mCH::H2B (magenta) (middle, bottom) from a male extruded gonad arm at the start (top, middle) and end (bottom) of a 90-minute image acquisition. Yellow brackets delineate the size of the spermatid region at the start of imaging. Scale bar = 20 µm. (D) Bright field images (top) and maximum intensity projections of SUN-1::GFP (inverted grey scale; bottom) from a late L4 hermaphrodite extruded gonad arm. Scale bar = 20 µm. Images on the right show a magnified view of the region boxed in the lefthand image. The dashed line indicates the same row of transition zone nuclei at the start (top) and after 2 hours of image acquisition (bottom). The arrowhead shows a germ cell that enters meiotic prophase during imaging. Scale bar = 10 µm. (E) Cropped timelapse images of germ cell nuclei at the distal (top) and proximal (bottom) ends of the distal gonad arm shown in (D). Arrowheads point to SUN-1::GFP foci appearing (top) and disappearing (bottom) over the course of imaging. Time is in minutes:seconds relative to the start of acquisition. Scale bar = 5 µm. (F) Single z-slices though a meiotic germ cell nucleus in a gonad explant from a late L4 hermaphrodite expressing SUN-1::GFP at the start (left) and end (right) of a 2-minute window during which SUN-1::GFP foci were tracked using TrackMate (middle). Images were acquired every 5 seconds. (G) Bright field image (top) and maximum intensity projections (middle, bottom) of CED-1::GFP from an adult hermaphrodite extruded gonad arm. The red arrowhead points to an engulfed apoptotic germ cell at the start of image acquisition (middle) and the yellow arrowheads indicate partially engulfed cells at the end of imaging (bottom). Scale bar = 20 µm.

While mitotic germ cell divisions were evident in the distal region of explants, we also observed spermatogenic meiotic divisions at the proximal end of explants from late larval hermaphrodites (Figure 1C; Movie 2 (Shakes et al., 2009)). To further assess the utility of explants for monitoring germ cell meiosis, we imaged gonad explants from late larval males and saw sustained meiotic divisions at the proximal end, consistent with ongoing spermatid production (Figure 2C; Movie 5). To monitor earlier stages of meiotic progression, we imaged gonad explants from late larval hermaphrodites bearing a GFP-tagged version of the LINC complex protein SUN-1 (SUN-1::GFP) (Machovina et al., 2016), which mediates chromosome movements in meiotic prophase and is required for homologous chromosome pairing and synapsis (Baudrimont et al., 2010; Penkner et al., 2009; Sato et al., 2009; Wynne et al., 2012). We observed that SUN-1::GFP levels were low in the distal mitotic region of gonad explants and increased in the so-called transition zone, where germ cells enter meiotic prophase (Figure 2D; Movie 6). Highly dynamic SUN-1::GFP puncta were observed in pachytene germ cell nuclei, over at least 120 minutes of imaging (Movie 6). We tracked SUN-1::GFP foci from one germ cell nucleus and found that the average velocity over two minutes in explants was strikingly similar to what has been observed in vivo (71±0.7 nm/sec compared to 76 nm/sec (Woglar et al., 2013); Figure 2E; Movie 7). Furthermore, we observed the appearance of dynamic SUN-1::GFP foci in distal transition zone nuclei, while foci disappeared progressively from proximal meiotic nuclei (Figure 2F; Movie 6), suggesting a sustained flux of germ cells through meiotic prophase over the course of image acquisition. Thus, germ cells properly progress through meiosis in gonad explants.

In hermaphrodite gonads, approximately half of all germ cells undergo physiological apoptosis, instead of differentiating into gametes (Chartier et al., 2021; Gumienny et al., 1999; Raiders et al., 2018). Germ cells enter apoptosis during meiosis, typically near the loop region of each gonad arm (the turn of the U), and are engulfed by somatic sheath cells via a CED-1-dependent pathway (Zhou et al., 2001). To assess whether germ cells undergo apoptosis in gonad explants, we performed live-cell imaging of adult hermaphrodite explants bearing GFP-tagged CED-1 (CED-1::GFP) expressed in the somatic sheath cells (Schumacher et al., 2005). We observed dynamic CED-1::GFP-positive protrusions throughout the loop region of gonad explants, as well as expanding pockets enriched for CED-1::GFP (Figure 2G; Movie 8), consistent with continuous germ cell apoptosis and engulfment.

Together, these results demonstrate that gonad explants can sustain germ cell mitosis, meiosis, apoptosis and gametogenesis, essentially recapitulating key properties that have been documented *in vivo*. Explants thus constitute a robust complementary model to characterize various dynamic processes in the germ line.

### *C. elegans* gonad explants are amenable to acute drug treatment

While conditional genetic perturbations, such as temperature-sensitive alleles or degron-mediated protein depletions, are more rapid than conventional alleles to probe gene function (Davies et al., 2017; Zhang et al., 2015), they often still take time to perturb gene activity and are rarely sufficient to interrogate dynamic events like mitosis, which in germ cells occurs on the order of minutes. The use of specific chemical inhibitors that rapidly target a protein of interest provides a valuable complement to these approaches. Such small molecule inhibitors have been used in *C. elegans*, yet the presence of the cuticle and efficient efflux transporters limits their effectiveness, especially in internal organs like the germ line (Roy, 2025; Xiong et al., 2017).

We used nocodazole to assess whether gonad explants are amenable to acute small molecule perturbations. Nocodazole is well characterized for its impact on microtubule dynamics and results in microtubule depolymerization in a wide range of cell types and species (Brabander et al., 1986; Florian and Mitchison, 2016; Vasquez et al., 1997); however, acute treatment of germ cells has only been achieved via microinjection (Kitagawa and Rose, 1999), thus limiting its usefulness as a tool. We compared the impact of nocodazole treatment on mitotic germ cells by measuring GFP::β-tubulin at germ cell centrosomes in gonad explants and intact animals treated with nocodazole during the mounting and imaging process. We measured centrosome GFP::β-tubulin at the start of image acquisition, approximately five minutes after nocodazole addition, and found that levels were unchanged in germ cells in intact animals but showed a dose-dependent decrease in germ cells in gonad explants (Figure 3).

**Figure 3.**
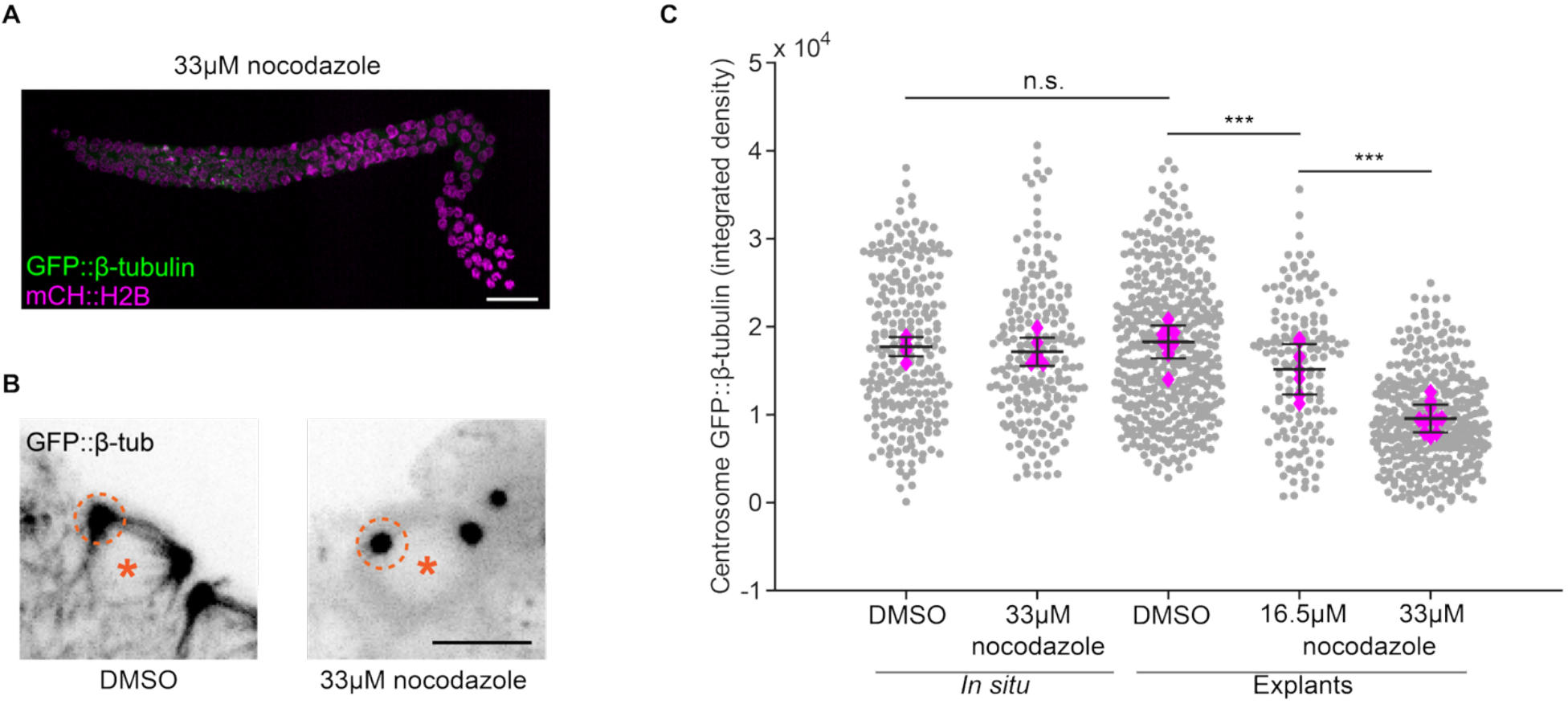
Gonad explants are amenable to acute nocodazole treatment. (A) Maximum intensity projection of GFP::β-tubulin (green) and mCH::H2B (magenta) from an extruded gonad arm treated with 33 µM of nocodazole. Scale bar = 10 µm. (B) Maximum intensity projections showing the change in GFP::β-tubulin in mitotic germ cells from gonad explants treated with DMSO (left) compared to 33 µM nocodazole (right). Both cells are in prophase. Spindle and astral microtubules are absent in the nocodazole treated cell. Dashed circles indicate the size and position of the regions of interest (ROIs) used to measure GFP::β-tubulin levels in (C). Asterisks mark the nuclei from which GFP::β-tubulin is largely excluded. Scale bar = 5 µm. (C) Integrated intensity of GFP::β-tubulin at the centrosomes of germ cells *in situ* and in gonad explants treated with DMSO or nocodazole, as indicated. Grey dots represent individual centrosomes and magenta diamonds represent the average measurement per gonad. Bars indicate average ± standard deviation. n.s. = not significant (p>0.05), *** = p<0.001, by an ANOVA with Tukey–Kramer post hoc test.

We next assessed the impact of nocodazole on chromosome congression and mitotic progression in germ cells. Microtubule depolymerization typically results in defects in chromosome congression and activates the SAC, blocking entry into anaphase and mitotic exit (Brabander et al., 1986; Lara-Gonzalez et al., 2021; Rieder and Maiato, 2004). Accordingly, we found that germ cells from nocodazole-treated explants showed aberrant mitotic figures (Figure 4A; Movie 4), with a significant number of germ cells showing chromosome congression defects, as assessed by the presence of more than one chromosome mass (Figure 4B). In addition, we found that gonad explants treated with nocodazole accumulated mitotically arrested germ cells over the course of image acquisition and this appeared to be dose-dependent (Figure 4C). Germ cell mitotic arrest following nocodazole treatment was dependent on the SAC, as arrested cells were not seen in explants carrying a null allele of the core SAC regulator and *MAD3* orthologue *san-1* (Figure 4C-D, Movie 9), despite significantly reduced GFP::β-tubulin levels at spindle poles (Figure 4E). Together, these results demonstrate that nocodazole treatment produces specific and acute effects in mitotic germ cells in *C. elegans* gonad explants.

**Figure 4.**
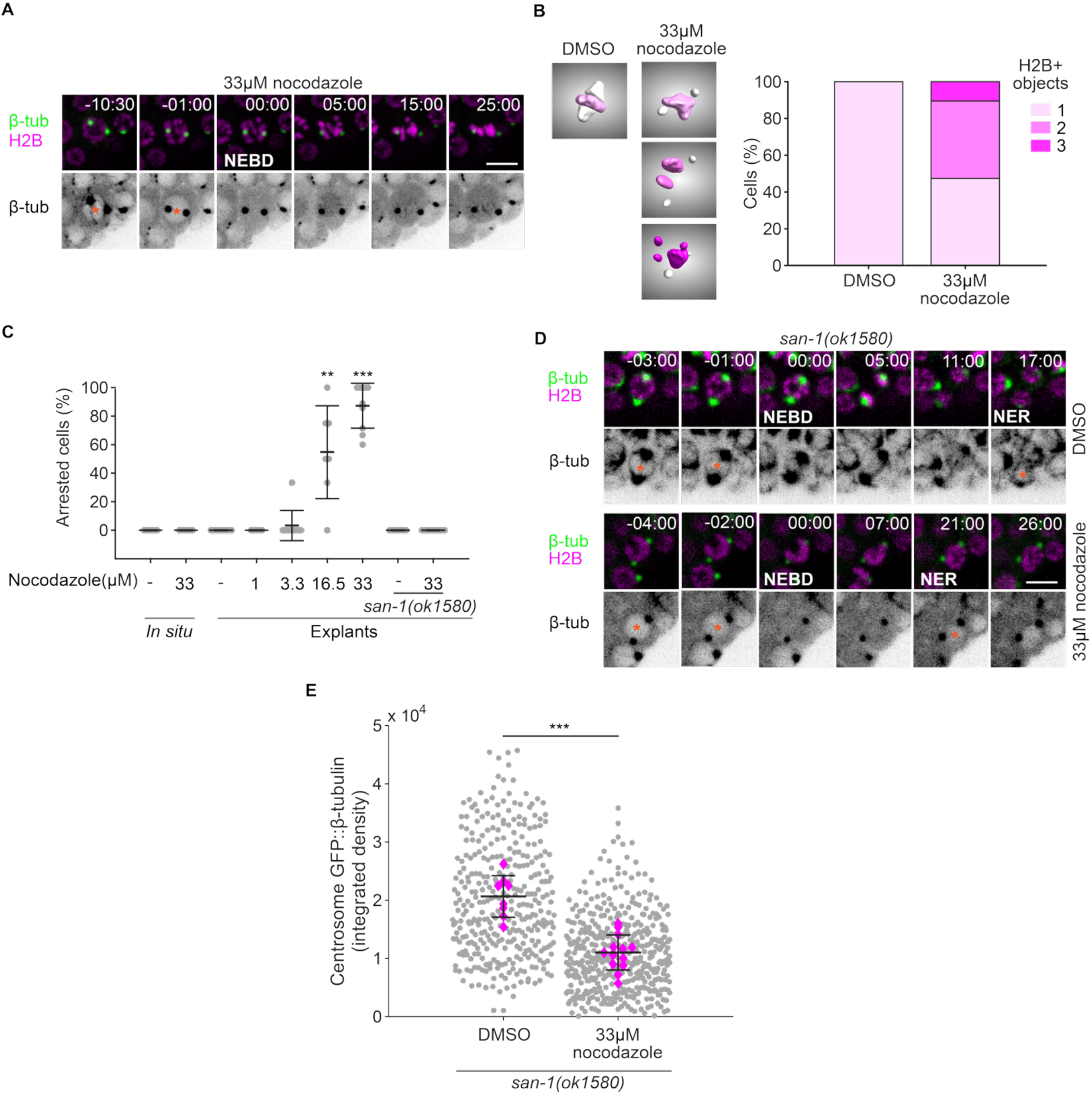
Nocodazole disrupts germ cell spindle assembly and leads to checkpoint-dependent mitotic delays. (A) Cropped timelapse images of an explant germ cell undergoing mitosis in the presence of 33 µM of nocodazole. The spindle is marked by GFP::β-tubulin (green) and chromosomes by mCH::H2B (magenta). Numbers indicate time in minutes:seconds relative to nuclear envelope breakdown (NEBD). Scale bar = 5 µm. (B) Three-dimensional surface renderings of GFP::β-tubulin (white) and mCH::H2B (magenta) in mitotic germ cells from gonad explants treated with DMSO (left) compared to 33 µM nocodazole (right). DMSO treated cells have one H2B-positive (H2B+) object at metaphase, while nocodazole treated cells often arrest with two or more H2B+ objects. The stacked bar plot shows the frequency of H2B+ rendered objects in both conditions. n = 19 cells in each condition. (C) Plot showing the percent of germ cells arrested in mitosis per gonad arm *in situ* and in gonad explants. Measurements in a checkpoint-null background (*san-1(ok1580)*) are shown on the right. Bars indicate average ± standard deviation. Each dot represents one gonad arm. ** = p<0.01, *** = p<0.001, by a Kruskal-Wallis with a Dunn post hoc test. (D) Cropped timelapse images of explant germ cells from *san-1(ok1580)* animals treated with DMSO (top) or 33 µM nocodazole (bottom). The spindle is marked by GFP::β-tubulin (green) and chromosomes by mCH::H2B (magenta). Numbers indicate time in minutes:seconds relative to NEBD. NER = nuclear envelop reformation. Asterisks mark the nuclei, as visualized by GFP::β-tubulin exclusion. Scale bar = 5 µm. (E) Integrated intensity of GFP::β-tubulin at the centrosomes of germ cells in gonad explants treated with DMSO or 33 µM nocodazole from *san-1(ok1580)* animals. Grey dots represent individual centrosomes and magenta diamonds represent the average measurement per gonad. Bars indicate average ± standard deviation. *** = p<0.001, by a two-sample t-test.

Altogether, this work establishes gonad explants as a new tool to characterize *C. elegans* germline function *ex vivo*. Germ cells in gonad explants retain normal mitotic and meiotic behaviors and are amenable to acute small molecule-induced perturbations. We note that while meiotic dynamics (SUN-1::GFP foci and spermatogenic divisions) appear relatively normal for up to two hours in explants, few new mitotic germ cells are seen after forty minutes of explant culture. This timing is consistent with what has been observed *in vivo*, where we and others have shown that germ cells rapidly arrest in the G2 phase of the cell cycle when animals are deprived of food (Seidel and Kimble, 2015; Zellag et al., 2021). Thus, explants may also provide a means to assess the signaling and/or metabolic factors underlying this response.

## Materials and methods

### Strain maintenance

*C. elegans* animals were maintained at 20°C on nematode growth medium (NGM; 2% agar, 3% peptone, 50 mM NaCl, 20 mM KH_2_PO_4_, 5 mM K_2_HPO_4_, 1 mM CaCl_2_, 1 mM MgSO_4_, 13 µM cholesterol) seeded with *Escherichia coli* (strain OP50) according to standard protocols (Brenner, 1974). Males and hermaphrodites of specific stages were selected based on the gonad and vulva morphology using a dissection microscope. The strains used in this study can be found in Table 1.

**Table 1.**
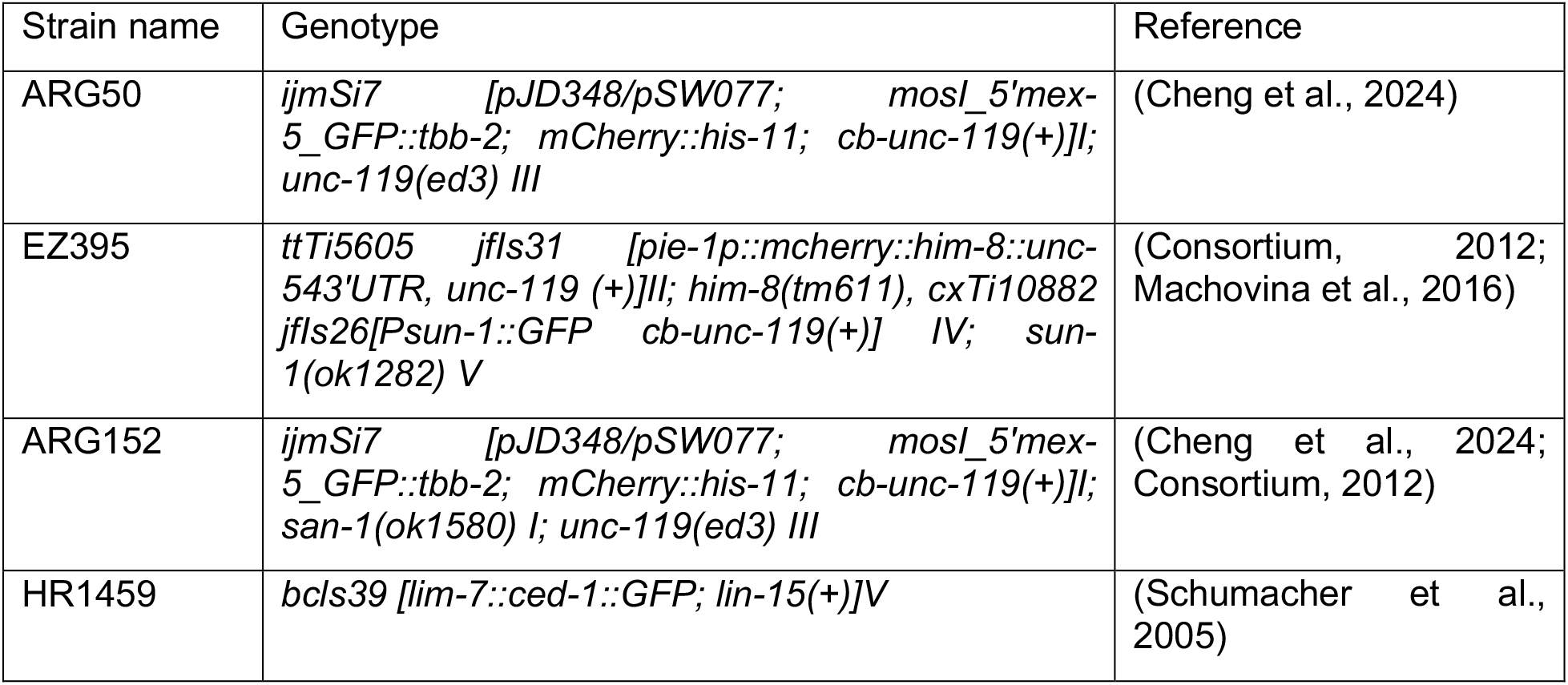
Strain list.

| Strain name | Genotype | Reference |
| --- | --- | --- |
| ARG50 | *ijmSi7 [pJD348/pSW077; mosl_5'mex-5_GFP::tbb-2; mCherry::his-11; cb-unc-119(+)]I; unc-119(ed3) III* | (Cheng et al., 2024) |
| EZ395 | *ttTi5605 jfls31 [pie-1p::mcherry::him-8::unc-543'UTR, unc-119 (+)]II; him-8(tm611), cxTi10882 jfls26[Psun-1::GFP cb-unc-119(+)] IV; sun-1(ok1282) V* | (Consortium, 2012; Machovina et al., 2016) |
| ARG152 | *ijmSi7 [pJD348/pSW077; mosl_5'mex-5_GFP::tbb-2; mCherry::his-11; cb-unc-119(+)]I; san-1(ok1580) I; unc-119(ed3) III* | (Cheng et al., 2024; Consortium, 2012) |
| HR1459 | *bcls39 [lim-7::ced-1::GFP; lin-15(+)]V* | (Schumacher et al., 2005) |

### Gonad extrusion and explant mounting

Using a mouth pipette and M9 buffer (22.04 mM KH_2_PO_4_, 42.27 mM Na_2_HPO_4_, 85.55 mM NaCl, 1 mM MgSO_4_), 10-15 animals were transferred from OP50-seeded NGM plates into the first well of a 3-well depression slide, filled with 100 μL of M9. Animals were swirled once to remove bacteria (as in (Zellag et al., 2022)) and transferred to the next well filled with 100 μL of M9 buffer plus 0.04% tetramisole (Sigma, Cat #L9756). After 1 minute anesthetization, animals were transferred to the third well filled with 100 μL of meiosis media (0.5 mg/mL Inulin (Sigma, Cat #I2255), 25 mM HEPES pH 7.5 (Wisent, Cat #600-032-CG), 60% Leibovitz’s L-15 Media (Wisent, Cat #323-050-CL), 20% heat-inactivated FBS (Wisent, Cat #090–150, lot 112740)) (Edgar and Goldstein, 2012), mixed once, as above, and transferred to a second, shallow depression slide filled with 50 μL of meiosis media. Gonads were extruded by sectioning the animals below the pharynx (Ananthaswamy et al., 2022; Lee and Schedl, 2006) using a single 25-gauge needle (BD Precision Glide #CABD305124) mounted on a 5 ml syringe for better grip. For adults, animals were cut at both the pharynx and the tail. Using a micropipette, 10 μL of meiosis media containing the extruded gonads, and some carcass material, was transferred to a glass slide patterned with 14 mm × 14 mm wells (Fisher Scientific, 30-2066A-BROWN 3 SQUARE 14 mm with Bars Epoxy autoclavable). A 22 mm × 22 mm cover glass was placed over the well and sealed completely with VaLaP (1:1:1 Vaseline, Lanolin, and Paraffin). Gonads that appeared undamaged and free from deformation by other tissues (see Figure S1) were selected for imaging. Note that meiosis media was filtered using a 0.2 µm syringe filter (Basix Cat#13100106) and stored at -20°C in 500 μL aliquots, which were re-filtered and maintained on ice prior to use.

**Figure S1.**
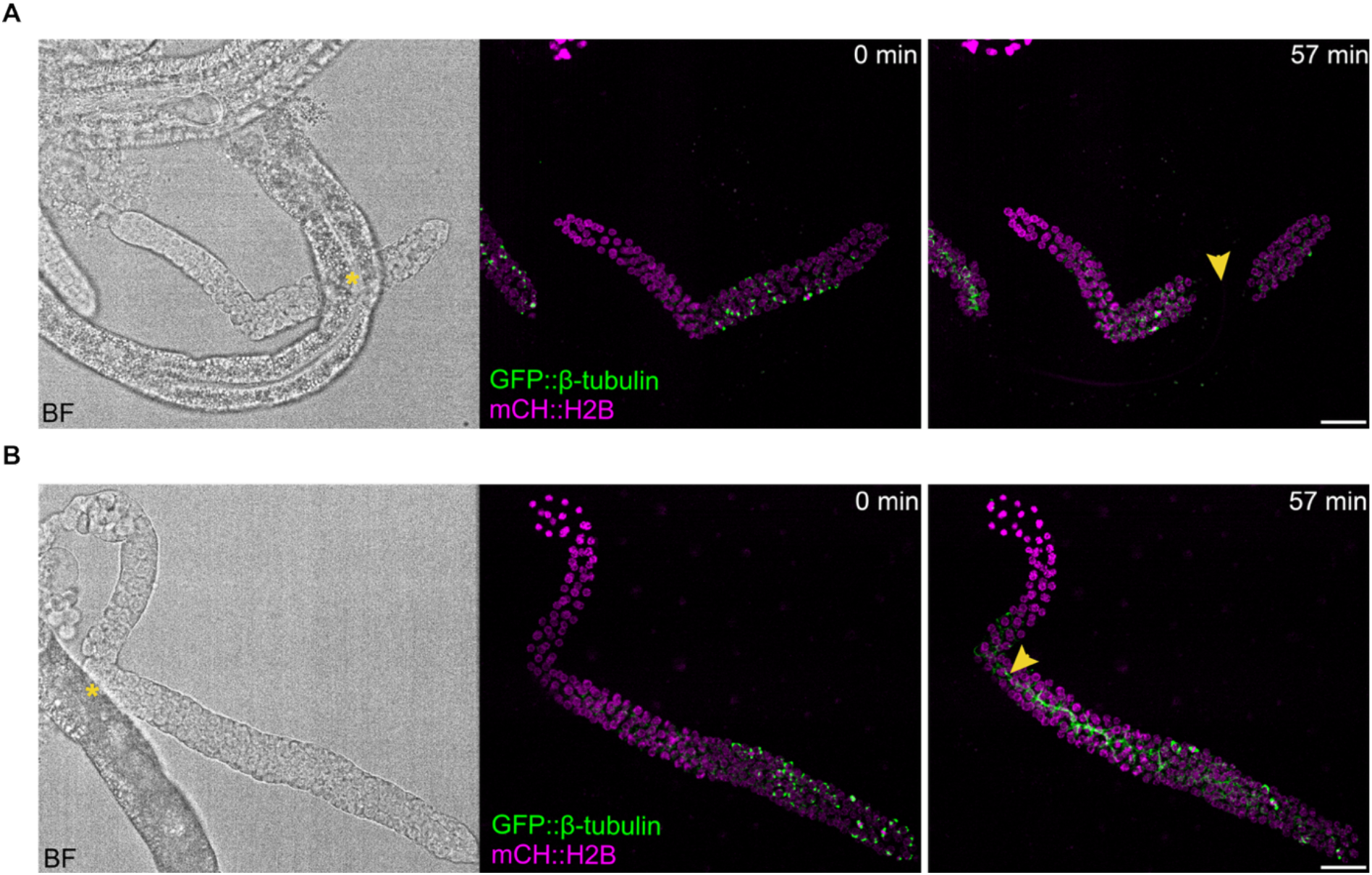
Examples of gonad explants that were not cleanly extruded and thus excluded from analysis. Maximum intensity projections of GFP::β-tubulin (green) and mCH::H2B (magenta) at the beginning (middle) and end (right) of the acquisition and bright field images at the beginning of acquisition (left). Gonad arms were extruded from L4 larval hermaphrodites. Asterisks show contact between the gonad explant and the gut. Arrowheads show aberrations that appeared during image acquisition. (A) Rupture of the gonad arm at site of contact with gut. (B) Fiber-like enrichment of GFP::β-tubulin at the presumed rachis surface suggesting a constricted or collapsed rachis. Scale bars = 20 µm.

### Whole worm *in situ* mounting

Whole worm *in situ* mounting was performed as described previously (Gerhold et al., 2015; Zellag et al., 2022; Zellag et al., 2021). Briefly, animals were transferred from OP50-seeded NGM plates using a mouth pipette and M9 buffer to a depression slide containing 100 μL of M9 buffer plus 0.04% tetramisole, before transferring to a 3% agarose pad, molded with grooves using a custom microfabricated silica plate. A 18 mm × 18 mm cover glass was placed, the corners were stabilized with VaLaP and the chamber was backfilled with M9 buffer plus 0.04% tetramisole before the edges were fully sealed with VaLaP.

### Nocodazole treatment

A 33 mM stock of nocodazole (Sigma, Cat #M1404) was prepared by solubilizing it in dimethyl sulfoxide (DMSO, Sigma, Cat #D4540). Final nocodazole concentrations were achieved by diluting this stock in either M9 buffer supplemented with 0.04% of tetramisole (for *in situ* mountings) or meiosis media (for explant mountings). Controls used a final concentration of 0.1% DMSO alone in the same buffers.

### Live-cell imaging

Images were acquired at room temperature (∼20°C) with a Nikon TI2-E inverted microscope controlled by NIS-Elements software and equipped with a Yokogawa CSU-X1 confocal head, using a Nikon Apo 40×/1.25 NA water immersion objective, 488 nm (100 mW) and 561 nm (100 mW) solid-state lasers, a dual band pass Chroma 59004m filter and a Hamamatsu ORCA-Fusion BT sCMOS camera. Acquisition setting were as follows: lasers at 5% intensity, 100 ms exposures, 0.5 μm z-step, every 1-3 min, for 40 to 120 min. Images of adult germline explants in Figure 1D were acquired using the same conditions except that a Nikon Plan Apo lambda 20x/0.75 NA objective and 1 μm z-step were used. Images of germline explants from animals expressing SUN-1::GFP (Figure 2D-F) were acquired using the same conditions except that a 1 μm z-step and a 200 ms exposure was used. For SUN-1::GFP foci tracking (Figure 2E) images were acquired every 5 sec. Images used for chromosome surface renderings (Figure 4B) were acquired using the same conditions except that a Nikon Plan Apo Lambda 60x/1.4 NA Oil immersion objective and 200 ms exposures were used.

### Measurement of mitotic duration

We used CentTracker, as described previously (Zellag et al., 2022; Zellag et al., 2021). Briefly, images were XY registered, centrosomes were tracked using TrackMate (Ershov et al., 2022; Tinevez et al., 2017) and automatically paired. The centrosome-to-centrosome distance (spindle length) overtime was used to define nuclear envelop breakdown (NEBD), prometaphase/metaphase and anaphase onset. The duration of mitosis was defined as the time between the end of NEBD (when spindle length reaches a minimum) and anaphase onset (when spindle length increases rapidly).

To score the number of germ cells per gonad that were arrested in mitosis, we counted the number of germ cells that underwent NEBD during the first 10 minutes imaging and that exited mitosis before the end of the acquisition versus the total number of germ cells that underwent NEBD during the first 10 minutes of imaging. To score NEBD and mitotic exit, we used centrosome/spindle dynamics, as described above, and the distribution of GFP::β-tubulin fluorescence, which is excluded from the nucleus prior to NEBD and after nuclear envelop reformation (NER) in telophase.

### Fluorescence intensity measurements of centrosome GFP::β-tubulin

We used Fiji software version 1.54r (Schindelin et al., 2012) to detect all centrosomes in the first frame, using Trackmate v7.14.0 (Ershov et al., 2022; Tinevez et al., 2017), with a 2.5 µm blob diameter and a manually adjusted detection threshold to maximize centrosome capture. Trackmate spots were exported as ROIs and used to measure the background-subtracted, integrated fluorescence intensity of GFP::β-tubulin for each centrosome. Background was measured for each gonad as the average fluorescence intensity within the nuclei of at least 20 non-mitotic germ cells. Note that these measurements include centrosomes from germ cells at different stages of mitosis, early prophase through telophase, and centrosome GFP::β-tubulin is correspondingly variable.

### Chromosome surface rendering

Timelapse acquisitions were processed using CentTracker and spindle midpoint coordinates were used to crop a 10 × 10 × 10 µm sub-stack for single mitotic germ cells. Single timepoints containing germ cells in metaphase (control) or arrested in prometaphase (nocodazole-treated) were imported into Dragonfly 3D World (Comet Technologies Canada Inc., Montréal, QC; version 2025.1). Background signal was excluded using intensity-based thresholding via the “multi-ROI from Multiple Otsu Splits” function. To exclude neighboring nuclei, the function “Separate Connected Component (26-connected)” was used, keeping only the objects within the target cell. A binary mask was generated using the “Create greyscale image from multi-ROI” function. The surface was rendered using the “Generate contour mesh” function. Final surface rendering was performed with “Remesh with Poisson surface reconstruction” using default smoothing parameters. The number of individual rendered objects per cell was counted manually.

### Quantification and statistical analysis

Statistical analysis was performed in MATLAB (MathWorks, Natick, MA; version 2024b). A two-tailed Student’s t-test (MATLAB ttest2) was used to compare the duration of mitosis between cells in explants versus *in situ*. An ANOVA with Tukey–Kramer post hoc test (MATLAB anova1 and multcompare) was used to compare centrosome fluorescence intensity measurements. A Kruskal–Wallis with Dunn post hoc test (MATLAB kruskalwallis and multcompare) was used to compare the percent of arrested cells. Adobe Illustrator (San Jose, CA; version 30.2.1) was used to assemble figures. Graphs were generated using MATLAB, saved as PDFs and imported into Adobe Illustrator to generate vector-based graphics in high resolution. All representative images were processed in Fiji version 1.54r to adjust brightness and contrast, add scale bars, crop and set pseudo-coloring. Processed images were exported as RGB TIFs (Schindelin et al., 2012). Final resizing and cropping was done in Adobe Illustrator.

## Acknowledgements

We thank Dr. Monique Zetka (McGill University, Montréal) for sharing the EZ395 strain and members of the Labbé, Gerhold and Weber labs and of the Québec Worm Group for stimulating discussions and advice. Some strains were provided by the *Caenorhabditis* Genetics Center, which is funded by the NIH Office of Research Infrastructure Programs (P40 OD010440). This work was funded by a grant from the Canadian Institutes of Health Research (PJT-525829) to ARG and JCL.

